# Integrated Bioinformatics analysis and Metabolomic Responses in finding novel therapeutic approach to treat viral myocarditis

**DOI:** 10.1101/2025.11.13.688209

**Authors:** Yao Lu, Zhou Qianlu, Wang Yujie, Shen Ailian, Zhou Xiaoxian, He Yuan, Wang Jing, Jiang Longwei, Yao Jianhua, Gu Rong, Zhao Jinxuan, Gao Xiangdong, Mu Dan

## Abstract

**Background:** Myocarditis is one of the most common health problems in young people. Despite imaging techniques and endomyocardial biopsies advancing, there is still inadequate for myocarditis characterization. Some studies have shown that <u>myocarditis</u> is closely associated with mitochondrial dysfunction. Therefore, this study was aimed to identify mitochondrial-related biomarkers and gene regulatory networks in myocarditis and potential therapeutic targets.

**Methods:** We downloaded GSE95368 dataset from GEO to get myocarditis gene expression profiles, then used EdgeR and AI algorithm for bioinformatics analysis of DEGs’ functions. Established CVB3-induced mouse myocarditis model via intraperitoneal injection, and assessed energy metabolism differences using targeted metabolomics. CVB3 stimulus induced myocarditis in H9C2 cells. We assessed mitochondrial dysfunction (ATP, MMP) with commercial kits, MAPK8 release via ELISA, and MAPK8, p-PI3K, p-AKT levels by western blot. Furthermore, a total of 8 patients primarily diagnosed with myocarditis were enrolled, and levels of MAPK8 and CK in serum were detected. SP600125, an inhibitor of MAPK8, was administrated to CVB3-infected mice to study its potential protective effect in viral myocarditis.

**Results:** We obtained 77 DEGs enriched in myocarditis. Analysis yielded 11 modules, MEpurple module genes linked to myocarditis progression were identified via WGCNA. MAPK8, NAMPT and ALB are associated with mitochondrial function. CVB3-infected mice showed cardiac inflammation and high MAPK8 expression in serum or heart tissues.The target metabolism results indicated altered central carbon metabolism distinguished CVB3 group from control, with higher D-Glucose-1-phosphate and D-Glucose-6-phosphate and lower L-Cystine, dCMP, IMP and Xylulose-5-phosphate levels. CVB3 treatment caused mitochondrial dysfunction in H9C2 cells (decreased ATP, MMP), and increased MAPK expression. Western blot showed MAPK8 consolidated the levels of p-PI3K and p-AKT. CK activity was notably higher in myocarditis patients than healthy individuals. MAPK8 levels in myocarditis patients serum also exceeded those in healthy individuals. Pearson analysis indicated that MAPK8 and CK may contribute to myocarditis progression via distinct mechanisms. In CVB3-infected mouse model with SP600125 treatment, cardiac inflammation and MAPK8, p-PI3k, p-AKT expression were reduced, confirming MAPK8’s crucial role in viral myocarditis.

**Conclusion:** Our study identified MAPK8 as a key biomarker of myocarditis, and it may be a potential therapeutic target for myocarditis.

## Introduction

Myocarditis is clinically described as an inflammatory process of cardiac myocytes. In developed countries, myocarditis is mostly caused by viral infectious pathogens (e.g., coxsackievirus, adenovirus). Many viruses can damage the heart both through direct viral processes or through indirect mechanisms associated with the host immune response(17). During the Coronavirus disease 2019 (COVID-19) pandemic, the incidence of myocarditis was steadily increasing(7). In addition, the epidemiologic studies also demonstrated that the SARS-CoV-2 infection increase the incidence of myocarditis at least 15-fold compared to patients that were not affected by SARS-CoV-2(21). Some patients may progress to dilated cardiomyopathy, which subsequently leads to heart failure. This is the most common malignant cardiovascular event among young people. Currently, cardiac magnetic resonance (CMR) imaging and immunohistochemical inflammation diagnostics have been utilized to diagnosis viral myocarditis. Despite the rapid development of new imaging techniques and endomyocardial biopsies, which remain inadequate in providing comprehensive characterization of the specific causative mechanisms of myocarditis as basis for a therapeutic decision(2,4). Therefore, establishing specificity diagnostic methods is crucial.

Mitochondrial are organelles that are able to adjust and respond to different stressors and metabolic needs within a cell, showcasing their plasticity and dynamic nature(20). Literature has revealed a fascinating correlation between the balance of energy demand and supply, and the structure of mitochondria(16). Current research suggests that mitochondria in myocardial cells continuously fuse and fission to produce sufficient ATP, thereby maintaining the sustained beating of myocardial cells(18). Among them, the oxidative respiratory chain plays a crucial role in maintaining the redox homeostasis and metabolic balance of myocardial cells(14). The redox homeostasis and the tricarboxylic acid cycle revolve around oxygen. Under normal physiological conditions, the energy substances and oxygen are in chemical equilibrium. When oxygen is insufficient, the organic molecule synthesis pathway is activation, synthesizing inflammatory mediators, cytokines, etc. In turn, mitochondrial dynamics orchestrates the cellular metabolism. The metabolic dysfunction, such as suppression of OXPHOS, decreased ATP synthesis and mtDNA depletion, which are involved in various pathological conditions(15). Previous reports have revealed that in myocarditis, oxidative respiratory chain dysfunction caused by impaired mitochondrial function occurs prior to myocardial tissue cell edema and necrosis(11).

Thus, we speculate that the abnormal intracellular energy metabolism resulting from impaired mitochondrial function in myocardial cells constitutes the molecular mechanism of viral myocarditis. Hence, identifying the target molecules underlying this mechanism holds the key to diagnosis of viral myocarditis. We will also discuss how optimizing energy metabolism may be used as an approach to prevent or treat myocarditis.

## Materials and Methods

### Artificial intelligence and Bioinformatics analysis

Search the disease dataset GSE95368 via the GEO database (https://www.ncbi.nlm.nih.gov/geo/). Conduct differential expression analysis on the data using the EdgeR package. Key genes were obtained using the artificial intelligence machine, learning algorithm Neuralnet. The workflow summarized the study design.

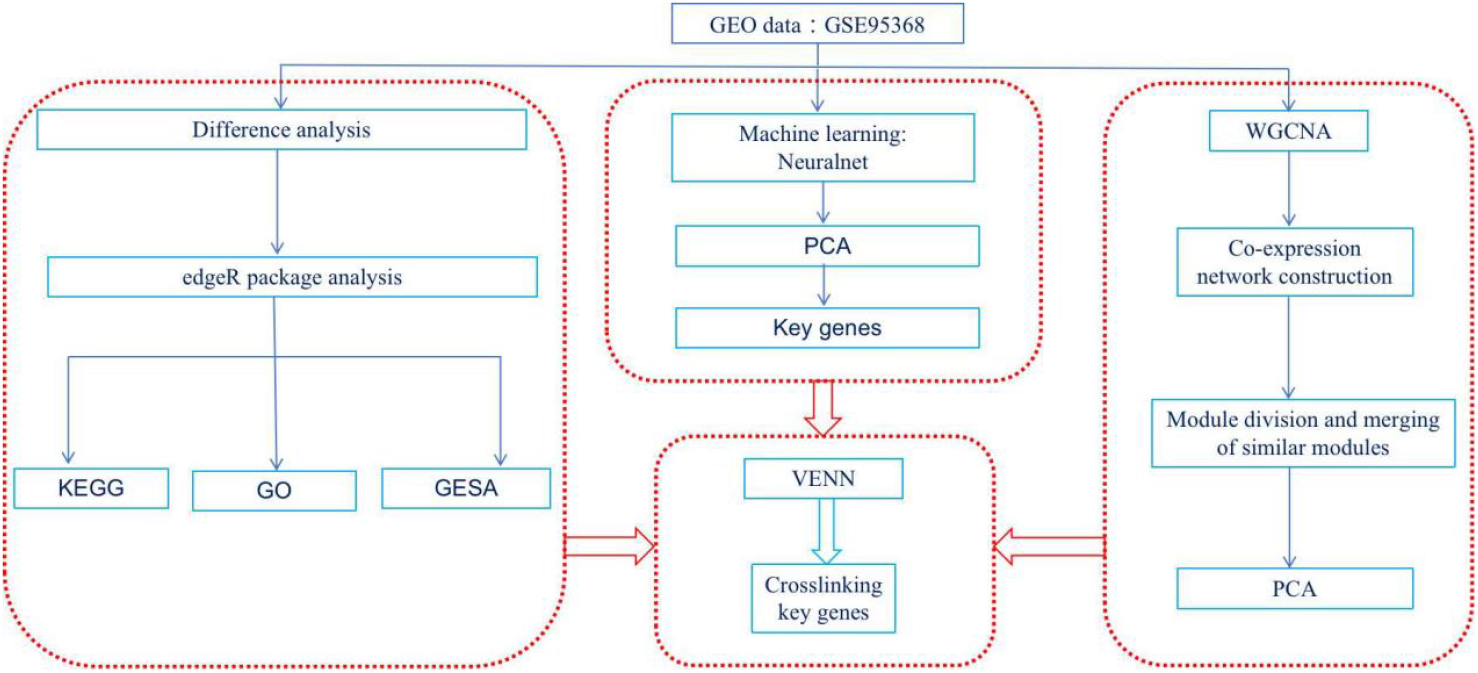

### Collection and statistics of clinical samples

Blood samples of 8 myocarditis patients initially hospitalized in the Nanjing Drum Tower Hospital were selected as the myocarditis group, and blood samples from 8 healthy people during the same period were selected as the control group. All patients were confirmed as myocarditis by imaging and pathology and had complete clinical data. Serum of all subjects were taken out of the refrigerator at −20°C, ELISA kit for MAPK8 detection (Elabscience) was performed according to the instruction of the ELISA kit and CK activity was detected by Amplex Red CK Assay Kit (Beyotime). The study was approved by institutional ethics committee Shanghai Tenth People’s Hospital following the Declaration of Helsinki (approval number: SHSY-IEC-6.0/25K75/P02). All parents/healthy people gave written informed consent. Using the concentration of standard substance as the horizontal coordinate and absorbance value as the vertical coordinate, plate a standard curve in order to calculate expression level of MAPK8 in serum.

### Experimental animals and models

All mice were bred in the Department of Laboratory Animal Science, Nanjing First Hospital in a standard specific pathogen-free(SPF) environment, and all animals were performed in accordance of the Ethical Committee of Nanjing First Hospital (approve number: DWSY-24048504). This study ensured all animal experimental procedures, including ethical principles, experimental design, operational standards, and regulatory frameworks, adhere to the guidelines from the NIH Guide for the Care and Use of Laboratory Animals. 4-5 weeks-aged male Balb/c mice were used (n=10 mice/group), and all required animal experimental rules were strictly observed. The mice were intraperitoneally inject with CVB3 (10^6^ TCID50, 200 μL), while mice in the control group were injected with the same amount of saline. In mice co-treated with SP600125, the inhibitor was injected irrigation (10 mg/kg), starting at 2 days before CVB3 infection, five days in all. CVB3 was donated from Zhongshan Hospital in Shanghai and virus titer was routinely determined by a 50% tissue culture infectious dose (TCID50) assay of H9C2 cell monolayer. Blood samples were obtained from the eye sockets of all mice, then the mice were euthanized by cervical dislocation to extract the heart tissue.

### Targeted metabolomics approaches

The targeted metabolomics analysis of central carbon metabolites using liquid chromatography(LC) combined with mass spectrometry(MS) in 6 cases of each group, to investigate of the metabolomics disorders of myocarditis. After the sample was thawed and smashed, an amount of 0.05 g of the sample was mixed with 500 µL of 70% methanol/water. The sample was vortexed for 3 min under the condition of 2500 r/min and centrifuged at 12000 r/min for 10 min at 4°C.Take 300 μL of supernatant into a new centrifuge tube and place the supernatant in −20°C refrigerator for 30 min, Then the supernatant was centrifuged again at 12000 r/min for 10 min at 4°C. After centrifugation, transfer 200 μL of supernatant through Protein Precipitation Plate for further LC-MS analysis. The sample extracts were analyzed using an LC-ESI-MS/MS system (Waters ACQUITY H-Class, https://www.waters.com/nextgen/us/en.html; MS, QTRAP® 6500+ System, https://sciex.com/). Linear ion trap (LIT) and triple quadrupole (QQQ) scans were acquired on a triple quadrupole-linear ion trap mass spectrometer (QTRAP), QTRAP® 6500+ LC-MS/MS System, equipped with an ESI Turbo Ion-Spray interface, operating in both positive and negative ion mode and controlled by Analyst 1.6.3 software (Sciex).

### Cells culture and treatment

H9C2 was obtained from the Cellcook Biotechnology Co., Ltd, China. The cells were cultured in a DMEM medium supplemented with (v/v) 10% FBS for 2 to 3 days until they reached 80% at 37°C in a humidified atmosphere of 5% CO_2_. The cells were treated with 100TCID50 CVB3 for 48 h. The supernatants were collected and cryopreserved immediately at −20°C for further experiments.

### Western blotting analysis

Total protein was extracted from the heart and H9C2 cells by RIPA lysis, then treated with loading buffer, denatured at 100°C for 10 min, and separated by gel electrophoresis. The transferred protein bands were blocked with 1% BSA and incubated in the following primary antibodies at 4°C overnight. The following primary antibodies were used: anti-MAPK8 (1:1000, Affinity Biosciences), anti-PI3K (1:1000, Cell Signaling Technology), anti-AKT (1:1000, Cell Signaling Technology), and anti-GAPDH (1:5000, Bioss). The membranes were then incubated with the following horseradish peroxidase-conjugated secondary antibody: HRP-labeled goat-rabbit IgG (1:10000, Beyotime). Immunoblot visualizations were performed using enhanced chemiluminescence. The relative expressions of the GAPDH were used to measure and assess the protein bands.

### Enzyme-linked immunosorbent assay

The supernatants from H9C2 cells were used to assess the concentrations of IL-6, MAPK8. ELISA kits (Eiaab Science) were performed in triplicates as per instructions of the manufacturer.

### Adenosine assay

Lysate from treated H9C2 cells were used to assess the adenosine concentration by fluorometric assay following the manufacturer’s instructions. Adenosine activity content was measured at 570 nm using a microplate reader, and the amount was normalized to the protein concentration.

### Creatine kinase activity assay

The serum was collected, and 20 μ L of the serum was used to measure creatine kinase activity according to manufacture’s instructions (Beyotime). To correct for background signal, the lowest value detected in CK assay buffer was subtracted from all other values. An enzyme activity unit (denoted as U) is capable of catalyzing the production of 1 μmol of ADP within a single minute at 37 °C.

### Mitochondrial membrane potential measurement

Mitochondrial membrane potential (MMP) was monitored by using JC-1 staining kit (Beyotime). After culturing, JC-1 was added to a final concentration of 2.5 μM at 37 °C for 15 min. Afterwards, MMP was visualized by fluorescence microscopy, and data were represented as the ratio of red to green fluorescence intensity.

### Histology, immunohistochemistry, and immunofluorescence staining

To assess the histopathological changes in heart mice, the paraffin embedded heart tissue was sectioned at 4 μm for HE staining, and the staining was performed following the manufacturer’s instructions.

For IHC staining, sections were used for staining of IL-6, TNF-α and MAPK8. For that, the heart tissue was hydrated, and non-specific antibody binds were prevented by flooding the slides with H_2_O_2_ for 10 min and then stopping with 5% BSA for 30 min. The slides were then incubated overnight with primary anti-IL-6 (1:200, Beyotime), anti-TNF-α (1:200, Beyotime) and anti-MAPK8 (1:200, Affinity Biosciences). The following day, tissues were incubated with goat anti-rabbit IgG and Streptavidin peroxidase for 25 min each, followed by DAB staining and hematoxylin counterstaining. The slides were dried in xylene and mounted for microscopy.

For IF, sections were used for staining of Ly6C/6G. For that, the heart tissue non-specific antibody binds were stopped with 1% BSA in PBS for 1 h. The slides were then incubated with primary anti-Ly6C/6G (1:200, Cell Signaling Technology) at 4 °C overnight, washed with PBS and probed with FITC-conjugated secondary antibody at room temperature for 1h. The slides were washed with PBS and incubated with DAPI nuclei staining for 10 min, followed by imaging.

### Small interfere RNA transfection

Downregulation of rat MAPK8 in H9C2 cells was achieved using following MAPK8 Rat Pre-designed siRNA from MedChemExpress. H9C2 suspension was transfected with siRNA-MAPK8 when plated following Lipofectamine RNA as the reverse transfection protocol.

### Lentivirus construction and transfection

To overexpress MAPK8 in H9C2 cells, the full-length cDNA sequence of rat MAPK8 was inserted into the lentivirus vector. H9C2 cells were infected with pLV-MAPK8 and the negative control in media without penicillin/ streptomycin or FBS for 6 h, and the media were then changed to culture media for 48 h before the next operation.

### Statistical analysis

Statistical analyses were carried out using GraphPad Prism 9. Statistical assays were performed as described in each Figure legend. Multiple samples were analyzed by one-way ANOVA test to evaluate statistical differences among the samples. The correlation between MAPK8 and CK of myocarditis patients was analyzed by χ^2^ test. Differences were considered statistically significant for *p* < 0.05.

### Data Availability Statement

The data generated in this study are available upon request from the corresponding author.

## Results

### 1 Screening Diagnostic Indicators for Myocarditis Utilizing Artificial Intelligence

AI biological analysis algorithms are proposed to construct artificial intelligence programs that perform deep analysis and learning on medical data, identifying potential disease-related targets within complex biological networks(24). Recent advancements in computational technologies have resulted in significant strides in applying AI in medical field. AI has been used to tackle various diseases, enhance imaging analysis, assist in nosology, and examine drug indication and therapeutic intent(10). With the utilization of AI in medical showing considerable promise, we use AI bioinformatics analysis to explore and investigate the mechanism by which impaired mitochondrial function leads to abnormal oxygen metabolism in viral myocarditis.

Using the edgeR package, we initially screened for 77 differentially expressed genes, comprising 51 upregulated genes and 26 downregulated genes. Subsequently, we employed the machine learning algorithm Neuralnet to identify 26 key genes (Fig.1A, 1B). Then, the KEGG database was utilized for the enrichment analysis of differentially expressed genes, while GO was employed for functional classification and enrichment analysis of these genes. It enriched them into important pathways such as MTOR, PI3K-Akt, MAPK, Wnt, Jak-STAT, EGFR, IL-17, and so on (Fig. 1C, 1D). Furthermore, the 26 key genes were collated into a gene set list, and the gene set enrichment analysis was by appropriate parameters and comparison conditions. The results showed that MAPK8, ALB, and LTA4H were enriched in the mTOR signaling pathway (Fig. 1E), thereby locking these genes and signaling pathways for further experimental verification.

**Figure 1.**
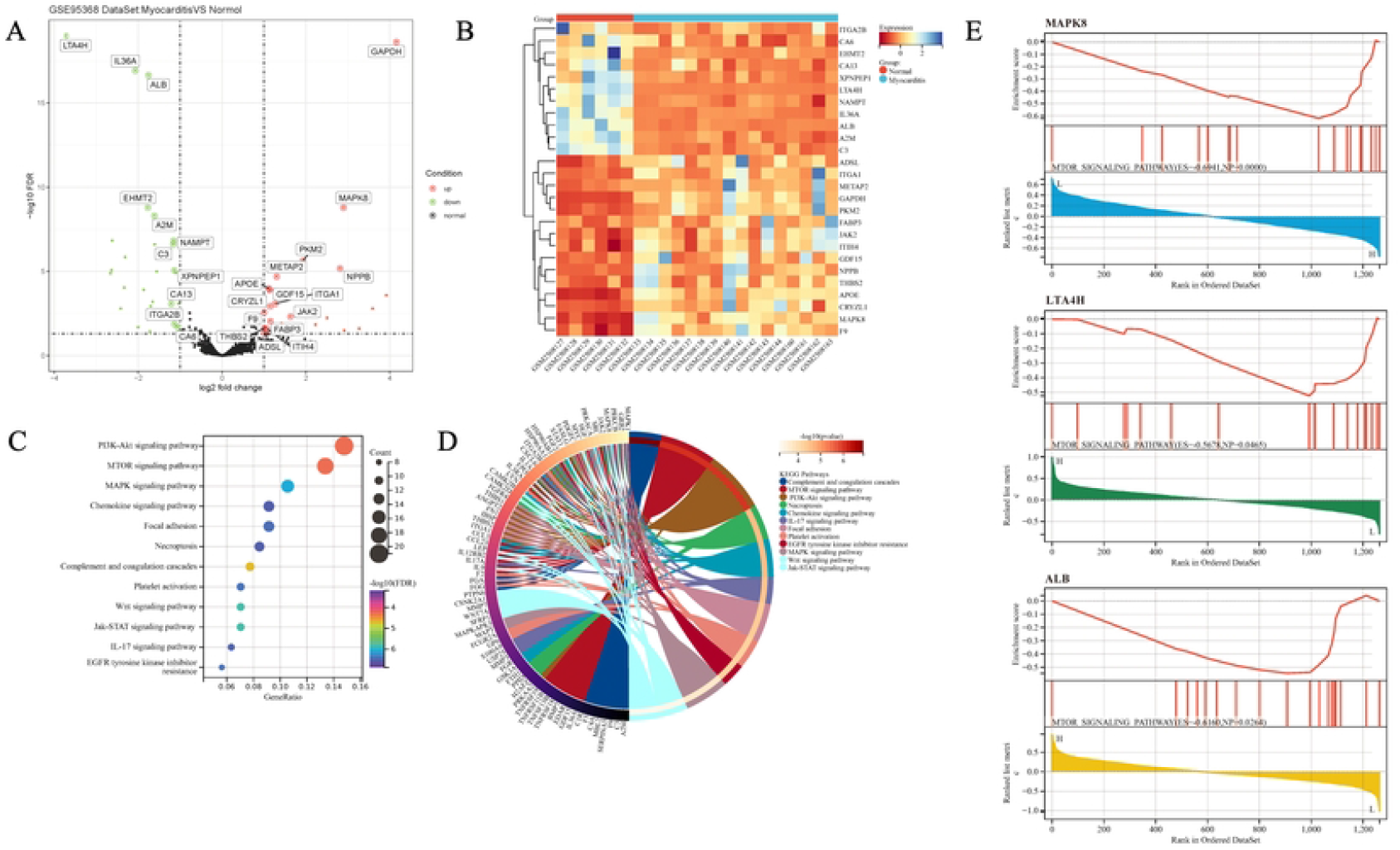

Based on the above results, Neuralnet, the Artificial intelligence computer algorithm, was utilized to conduct principal component analysis on the selected key genes, aiming to differentiate between the disease group and the normal group. The genes in the red circle represent the normal group, while the genes in the blue circle represent the myocarditis group (Figure. 2A). To explore the association between gene modules and phenotypes, we employed WGCNA (weighted gene co-expression network analysis) method to analyze the data and successfully divided the gene expression data into 11 modules (Fig. 2B), which were associated with different phenotypes. Among them, the MEpurple module genes exhibited a negative correlation with the progression of viral myocarditis, involving 46 genes, with a correlation coefficient of −0.95 and a p-value of 2E-11 (Fig. 2C). In addition, the high degree of differentiation between the MEpurple module gene and other modules indicates its significance in the myocarditis dataset, with a correlation coefficient of 0.93 and a P value of 9.6E-21(Fig. 2D). Both the heatmap and tree diagram results suggest the significance of the MEpurple module genes in the progression of myocarditis (Fig. 2E, 2F). Finally, the Venn map of genes obtained by WGCNA method, edgeR package differential analysis and AI machine learning, yielded 13 key genes, including LTA4H, ALB, EHMT2, A2M, NAMPT, C3, XPNPEP1, APOE, CA13, F9, CA6, MAPK8 (Fig. 2G). Beside, MAPK8, NAMPT, and ALB are associated with mitochondrial function and autophagy. Thus, we suggest that MAPK8 plays a significant role in the mitochondrial dysfunction involved in myocarditis.

**Figure 2.**
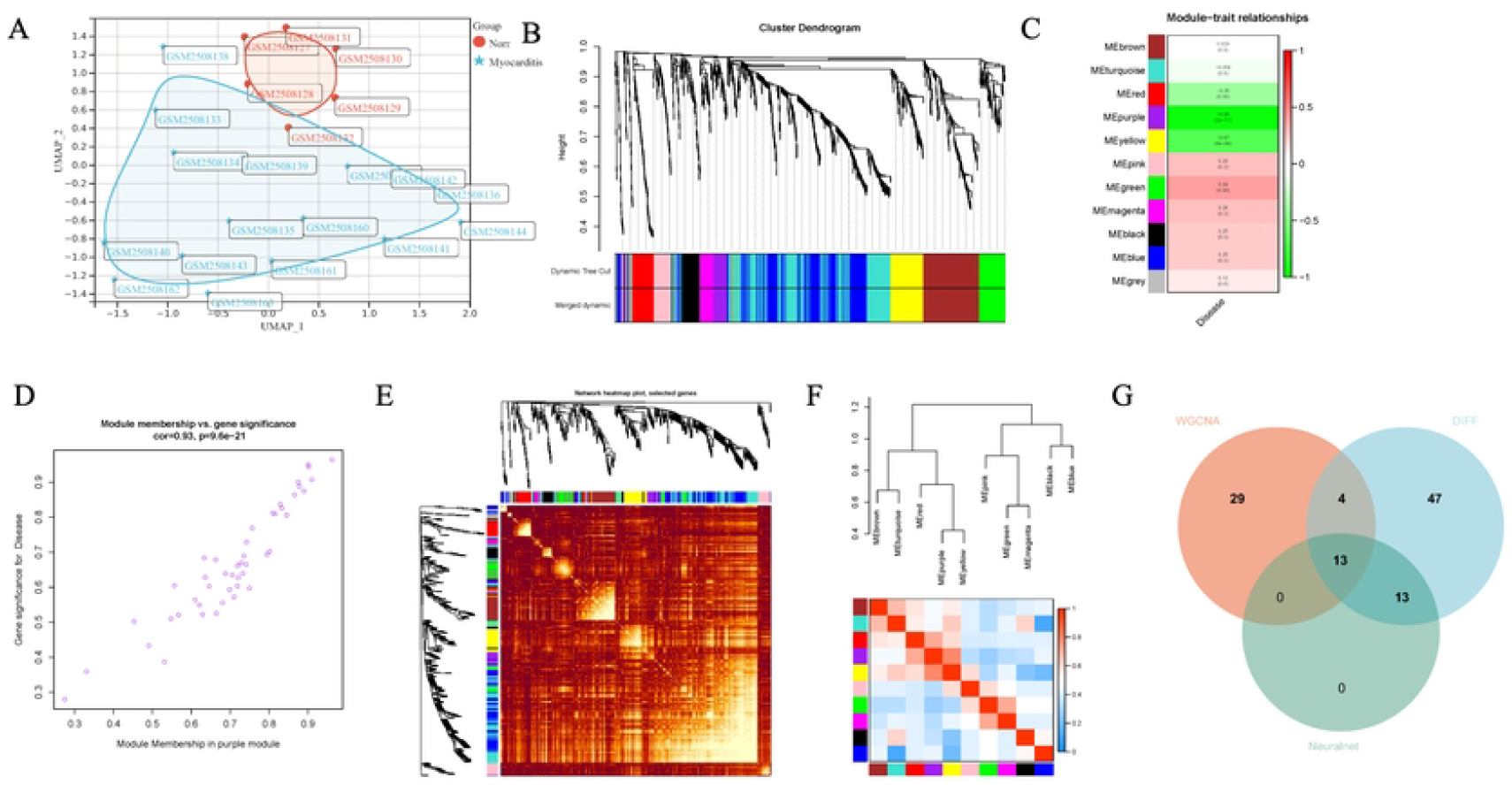

### 2 Assessment of differences in myocarditis mice by targeted metabolomics

Metabolomics uses advanced analytical chemistry techniques to enable the high-throughput characterization of metabolites from cells, organs, or tissues(1,19). The rapid growth in metabolomics is leading to a renewed interest in metabolism and the role that small molecule metabolites play in many biological processes. Metabolomics is yielding important new insights into a number of important biological and physiological processes. Thus, it has emerged as a valuable tool for identifying diagnostic biomarkers, unravelling molecular mechanisms and discovering potential therapeutic targets(20). To uncover key metabolic signatures and elucidate underlying mechanisms in viral myocarditis, we utilized metabolomics by targeting crucial metabolic pathways.

Mouse experimental models that utilized the Coxsackievirus group B type 3 (CVB3) virus infection causing myocarditis have illustrated the pathophysiology of viral myocarditis(27). Aaasys were carried out at 14 days after CVB3 infection(Fig. 3A). Results from HE staining indicated that significant inflammatory infiltration in the CVB3 group(Fig. 3B). Furthermore, the infiltration of Ly6C/6G-positive neutrophils induced by CVB3 infection were obvious(Fig. 3C). The level of serum MAPK8 was assessed using ELISA assay, and showed that MAPK8 was increased after CVB3 stimulation(Fig. 3D). Immunohistochemical density of IL-6, TNF-α were increased in the heart sections from mice treated with CVB3 compared to that from the control group(Fig. 3E). Moreover, the level of MAPK8 in heart sections exhibited a notable elevation compared to the control group, consistent with the result from ELISA.

**Figure 3.**
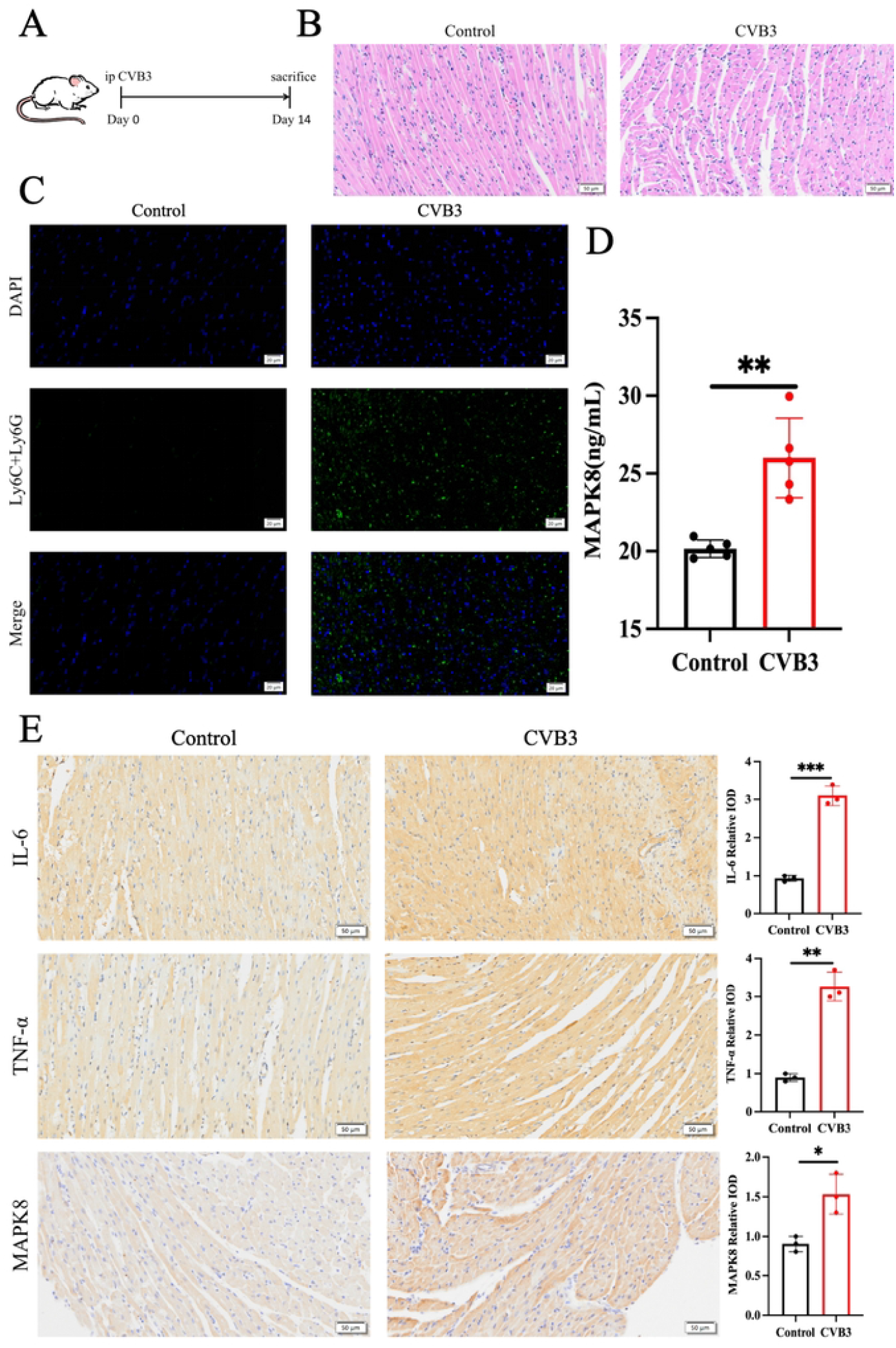

To analyze the metabolic changes between the CVB3 group and the control group, targeted metabolomics analysis was carried out using LC-MS. As a result, the PCA score plots showed a clear separation between the two groups(Fig. 4A). To further identify ion peaks that could possibly be used to differentiate the metabolic profiles of the two groups, the supervised OPLS-DA model was conducted. The result showed that the CVB3 group were also separated from the control group by the OPLS-DA score plots(Fig. 4B). To compare differences in metabolites between the two groups, differential metabolites were screened out based on VIP>1 in the OPLS-DA model and *P*<0.1 analyzed by Student’s *t*-test. The results showed that a total of 6 significant differential metabolites were identified between the two groups(Table. 1). The CVB3 group exhibited higher levels of D-Glucose-1-phosphate and D-Glucose-6-phosphate as well as lower levels of L-Cystine, dCMP, IMP and Xylulose-5-phosphate than the control group. These data confirmed that there were significant differences in central carbon metabolism between the two groups. Moreover, scatter loading plot analysis was carried out to evaluate whether these differential central carbon metabolites accounted for the distinguished profiles. The data suggested that altered central carbon metabolism could discriminate the CVB3 group from the control group(Fig. 4C). According to the metabolomics features, with the purpose of finding potential diagnostic biomarkers or therapeutic targets for myocarditis, we focused on the differential central carbon metabolites in CVB3 group compared to the control group. To facilitate the observation of patterns in metabolite content changes, we normalized the original content of the differential expressed metabolites obtained through screening on a row-by-row basis(Fig. 4D). To determine the pathway impact of these differential central carbon metabolites, we performed pathway classifications provided by KEGG analysis. Results indicated that the 6 differential central carbon metabolites were significantly enriched in 5 metabolic pathways, such as nucleotide metabolism, purine metabolism, carbon metabolism, amino acid biosynthesis, and cofactor biosynthesis(Fig. 4E, 4F). We subsequently evaluated the performance of these differential metabolites in distinguishing the CVB3 group from the control group. ROC analysis showed that, the central carbon metabolites classifier could act as potential disease diagnostic biomarkers in myocarditis(Table 2). In CVB3 group, the AUC values of two representative up regulated central carbon metabolites, D-Glucose-1-phosphate and D-Glucose-6-phosphate, respectively. Meanwhile, the ROC analysis also showed good diagnostic values of the down regulated central carbon metabolites, such as L-Cystine, dCMP, IMP and Xylulose-5-phosphate. The results above suggested that these key differential central carbon metabolites might be regarded as potential biomarkers for diagnosis or therapeutic targets of myocarditis. Besides, glycometabolism has been reported involved in PI3K/AKT/mTOR signaling during myocardial damage(5,22). However, it is unclear how MAPK8 and PI3K/AKT/mTOR signaling work together to damage mitochondrion via central carbon metabolism.

**Figure 4.**
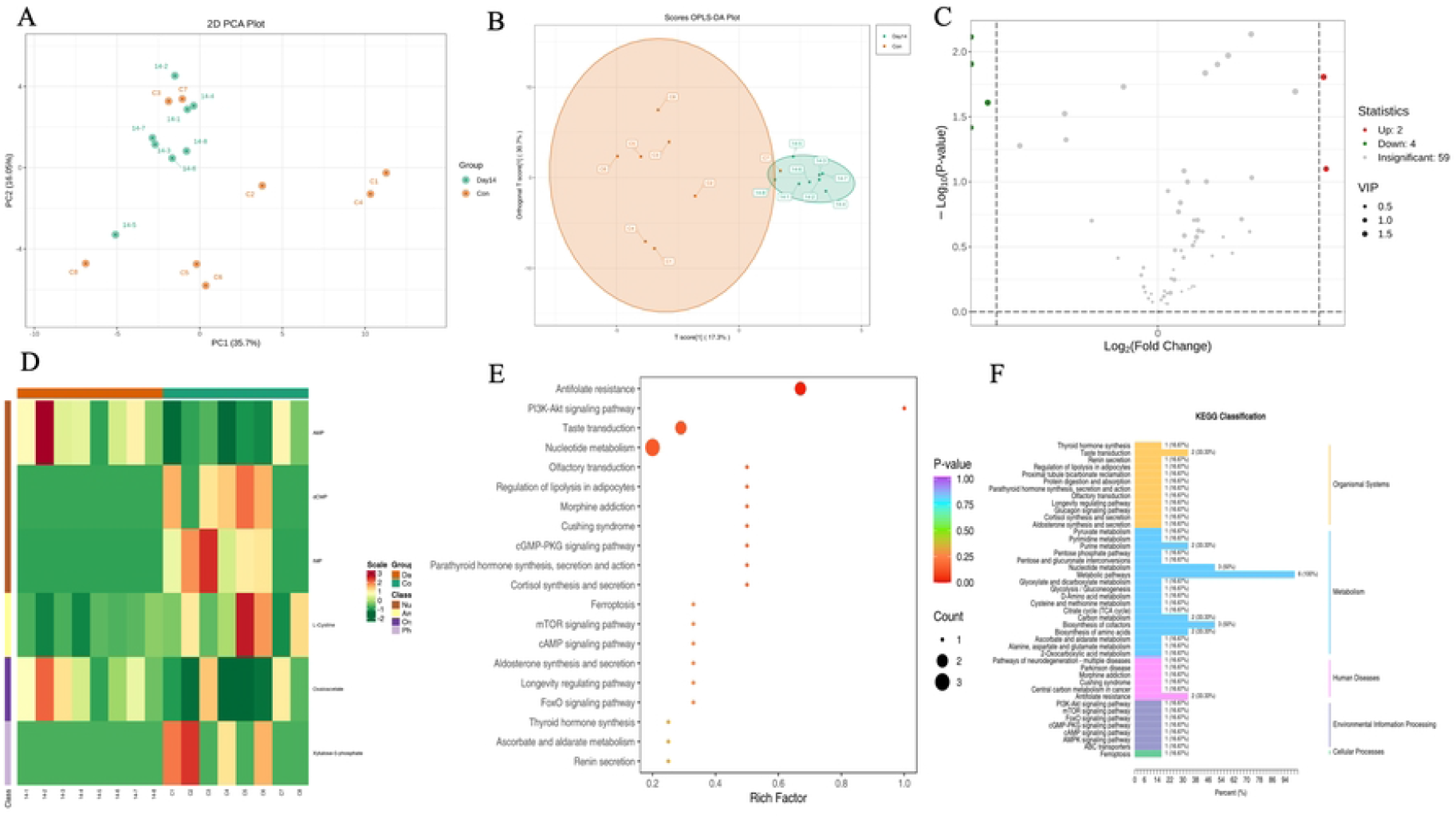

### 3 CVB3 induces mitochondrial dysfunction by disrupting the MAPK8/PI3K/AKT signaling pathway

In recent years, some studies shown that PI3K/AKT signaling play a key role in mediating cell survival and differentiation, proliferation, apoptosis, and metastasis(9,12). The PI3K/AKT signaling pathway is an important signal transduction bridge that connects extracellular signals with the signal transduction of cellular reactions that it plays a crucial regulatory role in various physiological functions of the cardiovascular system, including regulating the growth and survival of cardiomyocytes and the repair of cardiomyocytes after injury(8). However, there are currently no reports on myocarditis.

In this study, H9C2 cells were exposed to CVB3(100 TCID50) for 48 h, the secretion level of MAPK8 and IL-6 were notably elevated than the control group (Fig. 5A, 5B). mitochondrial plays a crucial role in cellular energy supply and participates in energy metabolism. To explore the effect of CVB3 on mitochondrial respiration function, we detected the level of intracellular ATP production. Fig. 5C displayed that CVB3 treatment stupendously weaken the basal respiration by decreasing the production of ATP. Changes in mitochondrial quantity can affect the normal function of cells. For the normal intracellular mitochondrial, the JC-1converted to its dye appeared in aggregate (red) form, and the JC-1 converted to its monomeric (green) form in the dysfunction mitochondrial(3,6). As demonstrated in Fig. 5D, the fragmented and dotted mitochondria emerged under CVB3 stimulus. To clarify the molecular mechanism of CVB3 damaging mitochondrial function, the expression levels of MAPK8-PI3K-AKT signaling pathway related proteins were detected by western blot technology. The protein expression level of MAPK8, p-PI3K and p-AKT were prominently increased under CVB3 stimulus (Fig. 5E). Additionally, the intervention of MAPK8 siRNA dramatically restrained the expression of MAPK8 and the phosphorylation level of PI3K and AKT (Fig. 5F). Simultaneously, the result in Fig. 5G disclosed that overexpression of MAPK8 by plasmid transfection overtly invigorate the phosphorylation level of PI3K and AKT.

**Figure 5.**
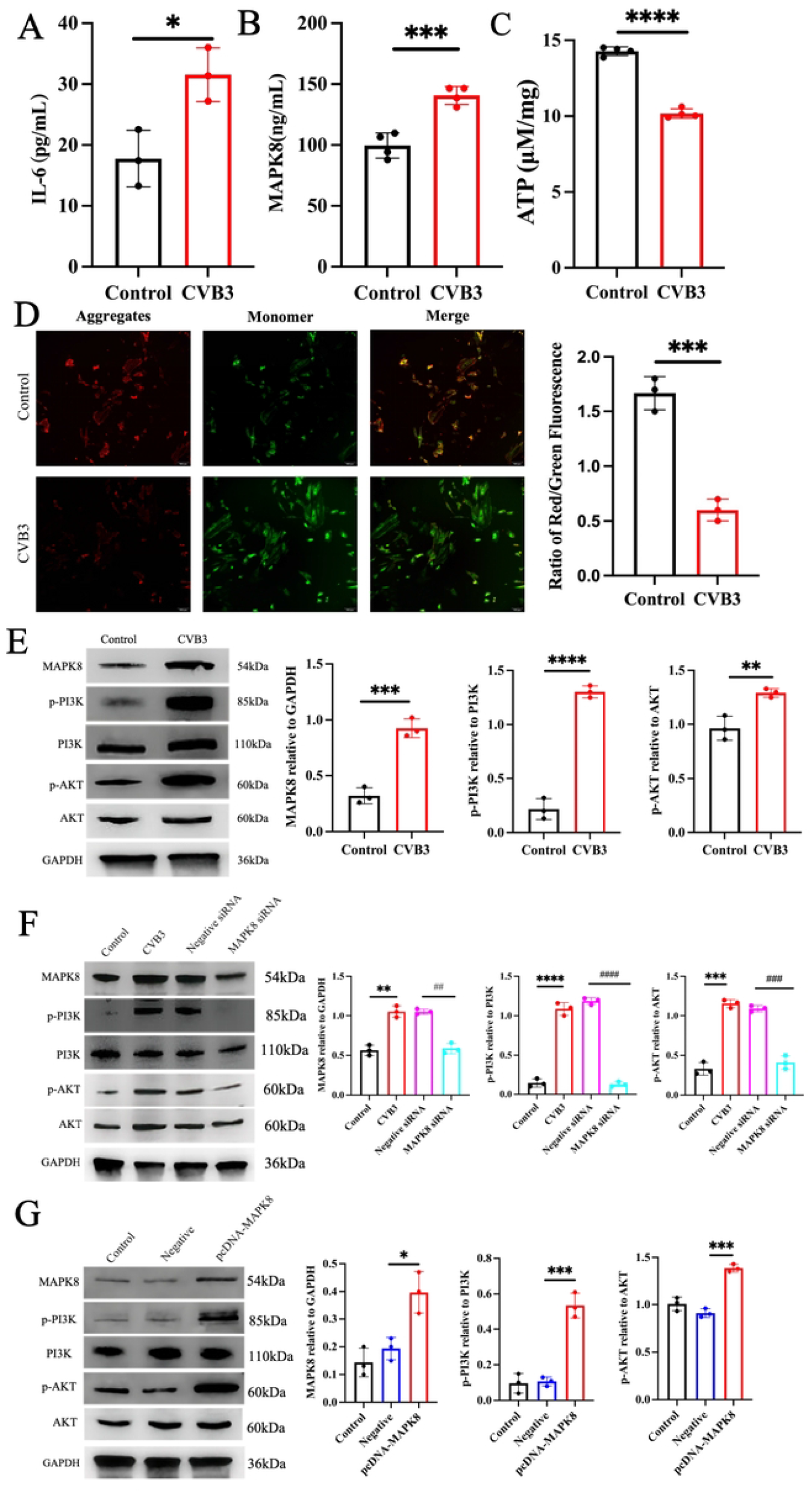

### 4 The serum of MAPK8 was notably elevated in myocarditis individuals

The expression levels of serum MAPK8 and CK in all groups were compared. The results showed that the expression levels of serum MAPK8 and CK in myocarditis patients were significantly upregulated compared to health controls (p<0.05) (Fig 6A, 6B). Meanwhile, the expression levels of serum MAPK8 and CK in myocarditis patients were poorly correlated (r=0.2192, *p*=0.4936) (Fig 6C, 6D). These results suggested that serum MAPK8 and CK levels in myocarditis patients are significantly higher than those in healthy people, and they may have different effects in the progression of the disease in myocarditis patients.

**Figure 6.**
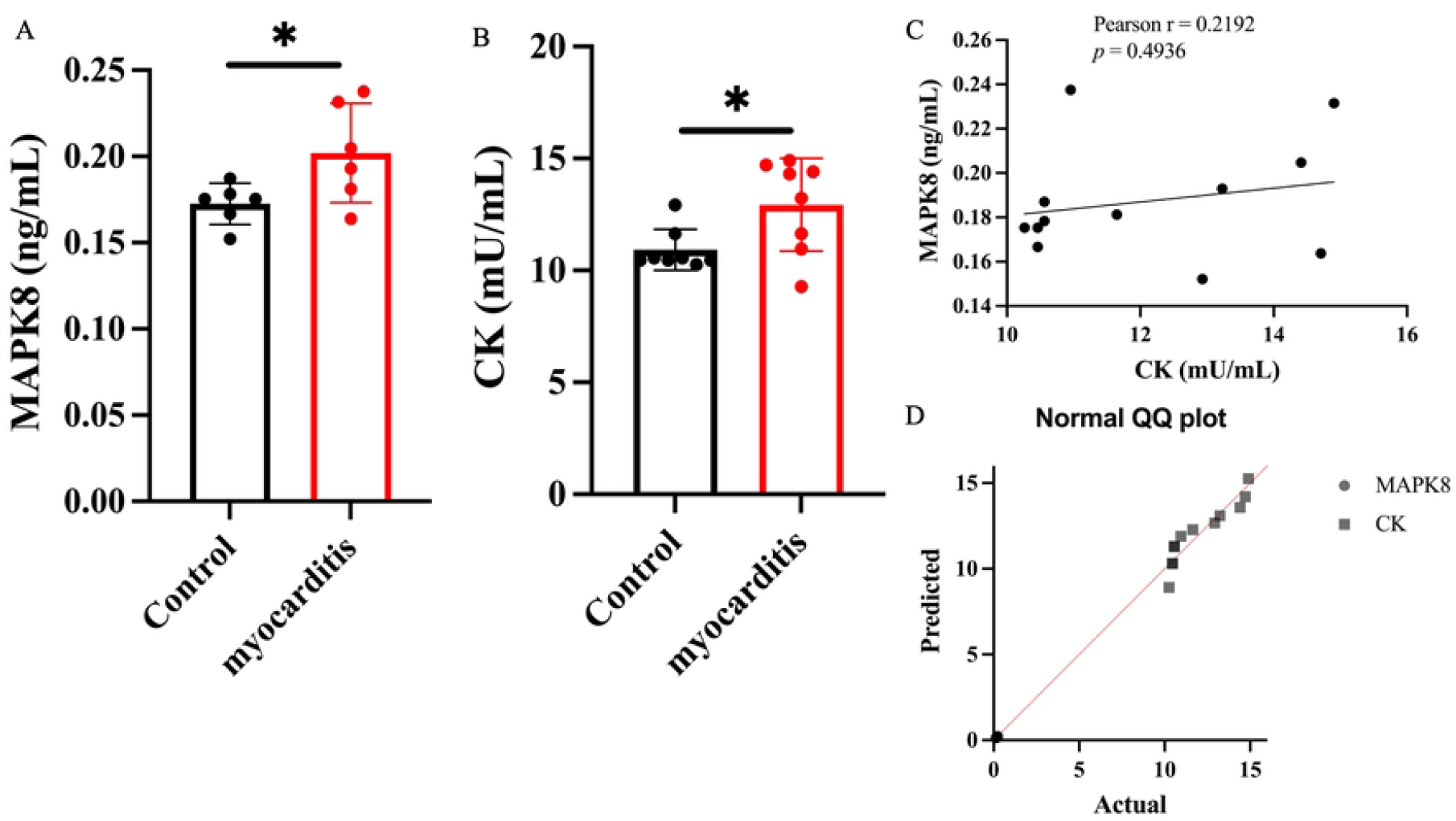

### 5 Protective effects of SP600125 on mice infected with CVB3

SP600125 is a reversible ATP competitive inhibitor of MAPK8 that is widely utilized in studies of MAPK8 pathway(23). In the present study, we investigated whether SP600125, as an inhibitor of MAPK8 signaling pathway, has potential value in clinical therapy of CVB3 infection *in vivo*.

To examine the protective effects of SP600125 on CVB3-infected mice, we administered SP600125 and Dexamethasone Sodium Phosphate (DEX) to mice respectively and observed the therapeutic effect (Fig 7A). Heart lesions were observed in mice on day 14 post-infection. There were dramatic differences in the histopathological changes in heart tissue between the CVB3 challenge and SP600125 treatment groups (Fig.7B). In the CVB3 challenge group, the heart tissue structure was loose and disorganized, accompanied by a significant infiltration of inflammatory cells. The pathological damage appeared to be less severe in mice treated with SP600125 or DEX, in which only slight inflammatory cellular infiltration. There were no obvious lesions in the heart of the mice in the control group. The result indicate that SP600125 treatment can effectively reduce the severity of pathological lesions in the hearts of CVB3-infected mice. To investigate the mechanism by which SP600125 inhibits infection, we measured the secretion level of MAPK8 by ELISA in the serum (Fig. 7C). The level of MAPK8 in the serum of SP600125-treated mice was evidently lower those in untreated mice on day 14 postinfection. Moreover, the expression of IL-6, TNF-α, and MAPK8 in the hearts were detected by ELISA (Fig. 7D) or immunohistochemistry (Fig. 7E). As shown in Fig. 7D and Fig. 7E, the expression levels of IL-6, TNF-α and MAPK8 in the hearts of the SP600126 treatment group mice were lower than those in the CVB3 challenge group, and the expression levels differed significantly. In addition, the expression of MAPK8, p-PI3K and p-AKT were detected by western blot in the hearts (Fig. 7F). The levels of MAPK8, p-PI3K, p-AKT expression in the hearts of the CVB3 challenge group were higher than in the SP600125 treatment group on day 14 postinfection. These data were consistent with the heart histopathologic changes described above. Our results suggest that SP600125 treatment can significantly reduce the inflammatory response in the hearts of CVB3-infected mice and downregulate the expression of MAPK8.

**Figure 7.**
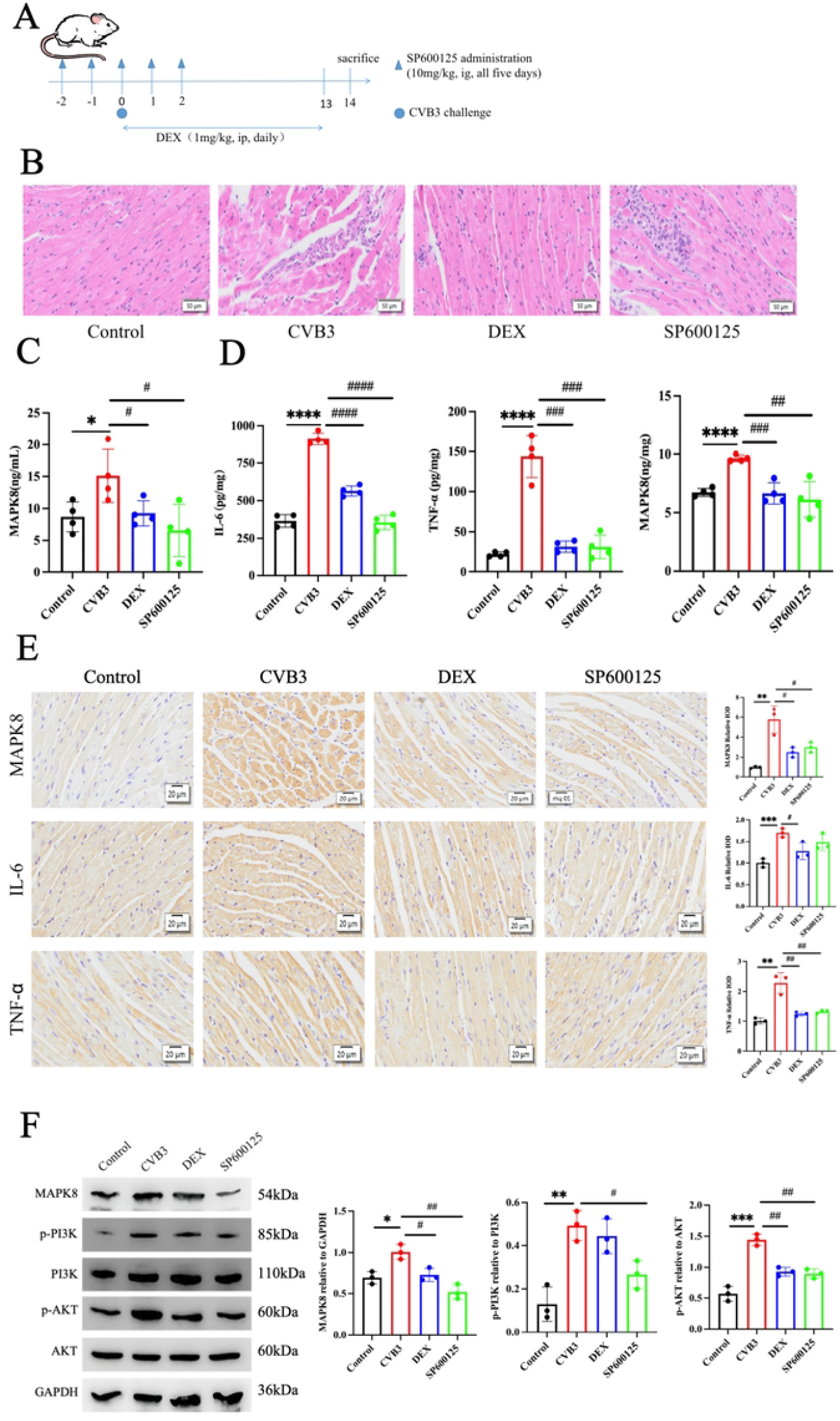

To determine the impact on cardiac energy metabolism of the treatment with SP600125, we performed energy metabolomices analysis. After CVB3 infection, acquisition of cardiac increased the levels of metabolites involved in diverse metabolic pathways, including the amino sugar and nucleotide sugar metabolism, biosynthesis of nucleotide sugars as well as biosynthesis of cofactors, as shown by heatmap and metabolic pathway enrichment analysis representations (Fig. 8A, 8B). Important differences were indeed found in the specific metabolites, notably D-Fructose-6-phosphate, D-Mannose-6-phosphate, D-Glucose-6-phosphate, D-Glucose-1-phosphate and D(+)-Glucose (Fig. 8C). Clearly, when CVB3-infected mice with SP60025 treatment, these metabolic changes were ameliorated. Several studies demonstrated that during myocarditis, the body may be in a state of stress, which can lead to sympathetic nervous system activation, thereby affecting glucose metabolism and causing stress hyperglycemia(28). Our results indicated that SP600125 could enhance myocardial mitochondrial function, thereby influencing myocardial energy metabolism.

**Figure 8.**
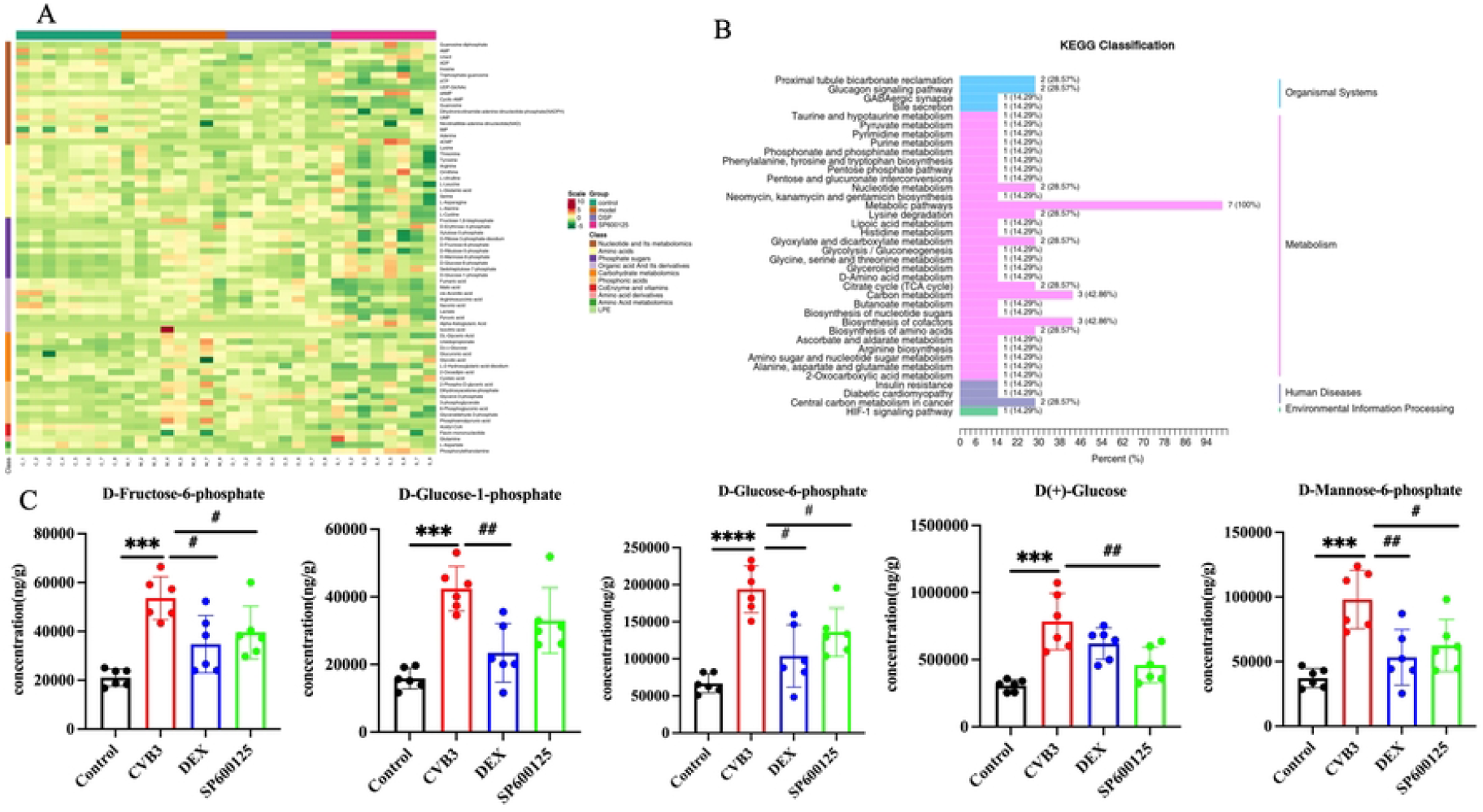

## Discussion

Viral myocarditis has an incidence of 10 to 22 per 100,000 individuals. The presentation pattern of viral myocarditis can range from nonspecific symptoms to more aggressive symptoms. After the initial acute phase presentation of viral myocarditis, the virus may be cleared or the viral infection may persist. The viral infection may lead to a persistent autoimmune-mediated inflammatory process with continuing symptoms of heart failure. As a result of these possibilities, the diagnosis, prognosis and treatment of viral myocarditis can be extremely unpredictable and challenging for the clinician(2). Viral myocarditis inflicts significant harm on patients physical and mental health, while also posing a considerable burden on society. Hence, research into its pathogenesis and drug treatment targets is of utmost importance.

Recent studies have shown that artificial intelligence technique is powerful tool in the discovery of novel target. Our study utilized myocarditis gene expression microarray data from the GEO public database. Through bioinformatics analysis methods and AI machine learning, we identified 13 key genes, namely LTA4H, ALB, EHMT2, A2M, NAMPT, C3, XPNPEP1, APOE, CA13, F9, CA6 and MAPK8, which are highly correlated with myocarditis. Wherein, LTA4H, NAMPT and MAPK8 are associated with mitochondrial function which were enriched in the PI3K/AKT signaling pathway. This provides new insights for the diagnosis and treatment of myocarditis.

Metabolomics is an omics technology that has emerged following genomics, transcriptomics, and proteomics. It enables the qualitative and quantitative analysis of all low molecular weight metabolites within an organism. Small molecule metabolites, as the final products of gene expression, can directly and accurately reflect the body’s pathological and physiological state(25). The application of metabolomics in myocarditis is increasingly widespread, offering more avenues for discovering new targets in the treatment of myocarditis. In recent years, the potential link between mitochondrial function and myocarditis has garnered widespread attention from researchers. This study established an animal model of viral myocarditis through CVB3 infection, and assessed the effectiveness of the model by examining inflammatory infiltration and inflammatory cytokines in cardiac tissues. Virus-infected mouse heart tissues exhibited pronounced inflammatory infiltration and increased expression levels of inflammatory cytokines (IL-6, TNF-α). Additionally, the level of MAPK8 significantly increased in both serum and heart tissue, which was consistent with the AI prediction results. Based on this, we conducted a qualitative analysis of the heart tissues of mice in the normal control group and CVB3-infection group using targeted metabolomics technology, identifying differential energy metabolites and characterizing six metabolites with higher levels of D-Glucose-1-phosphate and D-Glucose-6-phosphate as well as lower levels of L-Cystine, dCMP, IMP and Xylulose-5-phosphate. We observed differential energy metabolism between CVB3 group cardiac and control cardiac, establishing its significant associations with prognosis in myocarditis.

To clarify the molecular mechanism of MAPK8 in viral myocarditis, we treated H9C2 cells with CVB3 stimulus. As shown, the mitochondrial function assessment announced that CVB3 decreased the production of ATP. Coincidentally, under CVB3 stimulus, the decreased MMP was evidenced by increased green/red fluorescence signal by JC-1 staining. The pro-survival kinase-signaling cascade phosphatidylinositol 3-kinase/protein kinase B (PI3K-AKT) pathway is an intracellular signal transduction pathway to exert a pivotal role in cardiovascular system. Previous bioinformatics analysis also showed that MAPK8 is associated with mitochondrial function which was enriched in the PI3K/AKT signaling pathway. Thus, further investigation of the PI3K/AKT signaling pathway may provide new insights for the potential targets for treating myocarditis. The expression levels of PI3K-AKT signaling pathway related proteins were detected by western blot technology. The protein expression level of MAPK8, p-PI3K and p-AKT were prominently elevated under CVB3 stimulus. Additionally, the results disclosed that overexpression of MAPK8 increased the phosphorylation level of PI3K and AKT. Further evidence suggested that the intervention of MAPK8 siRNA dramatically restrained the phosphorylation level of PI3K and AKT. Notably, our results demonstrated that CVB3 impair mitochondrial function by prominently activating MAPK8-PI3K-AKT signaling pathway.

To further confirm the significance of MAPK8 in myocarditis patients, we analyzed the serum level of MAPK8 via ELISA kits. Unsurprising, the level of serum MAPK8 is upregulated in myocarditis individuals, which was consistent with above results. To explore the relationship between MAPK8 and serum creatine kinase (CK) level in myocarditis patients, we established the relationship using spearman analysis. As shown, there is a poorly relationship between serum level of MAPK8 and CK in myocarditis patients serum, they may exert an influence on the development of myocarditis via distinct pathways.

Aberrant kinase activity is associated with CVB3 infection, but no small-molecule kinase inhibitors are being tested in clinical of CVB3 infection(13). MAPK8 is biologically active and widely distributed in cells and tissues. Therefore, MAPK8 is presumed to be potential treatment target for various diseases(26). In this study, we explored the protective role of the MAPK8 inhibitor SP600125 in myocarditis induced by CVB3 infection in a mouse model. In the present study, we showed that SP600125 could decrease the severity of illness, and reduce the inflammatory response in the heart of CVB3-infected mice. CVB3 infection can result in a “cytokine storm”, in which high expression levels of inflammatory cytokines recruited to the infection site to trigger viral myocarditis. Our data showed that the expression of IL-6 and TNF-α was elevated on day 14 post-infection. However, SP600125 treatment led to significant downregulation of these cytokines, suggesting that inhibition of MAPK8 may have a strong influence on cytokine expression. Moreover, western blot results manifested that the expression of MAPK8, p-PI3K and p-AKT proteins in CVB3-infected mice was memorably elevated, while SP600125 treatment reversed this trend. Our data conclude that SP600125 treatment targeting the MAPK8/PI3K/AKT pathway may provide a novel antiviral treatment option against CVB3 infection.

After being administered with SP600125, the experimental mice exhibited an array of remarkable alterations in their energy metabolism pathways which were different from the CVB3-infected group. Glucose is an important source of fuel for the heart. The CVB3-infected heart undergoes dramatic alterations in energy substrate utilization and overall metabolic profile. Important differences were indeed found in the specific metabolites, notably D-Glucose-6-phosphate and D-Glucose-1-phosphate. While, when CVB3-infected mice were treated with SP60025, the metabolic disturbances that had arisen due to the viral infection were notably ameliorated. The therapeutic intervention with SP60025, a compound with known modulatory effects, seemed to exert a beneficial influence on the metabolic pathways disrupted by the CVB3 infection. Above results indicated that the administration of SP600125 to mice with viral myocarditis not only mitigated the inflammatory response and myocardial damage by modulating the MAPK8-PI3K-AKT signaling pathway but also markedly facilitated the remodeling of their energy metabolism mechanisms. This, in turn, provided cardiomyocytes with more ample energy support, representing a pivotal aspect of the drug’s therapeutic efficacy.

In summary, our data suggested that MAPK8 may serve as a potential biomarker of viral myocarditis. Mechanistically, the activated MAPK8-PI3K-AKT pathway by CVB3 may inhibit the mitochondrial function following energy metabolism abnormalities.

## Author contributions

Conceptualization and writing—original draft: Yao Lu; Data collection, Formal analysis: Zhou Qianru, Wang Yujie; Literature search: Shen Ailian, Zhou Xiaoxian, He Yuan, Wang Jing; Supervision, Funding Acquisition: Jiang Longwei, Yao Jianhua, Gu Rong, Zhao Jinxuan; Review and editing: Gao Xiangdong, Mu Dan. All authors approved the final version for publication.

## Ethical approval

All applicable international, national, and / or institutional guidelines for the care and use of animals were followed.

